# *C. elegans* epicuticlins define specific compartments in the apical extracellular matrix and function in wound repair

**DOI:** 10.1101/2024.01.12.575393

**Authors:** Murugesan Pooranachithra, Erin M. Jyo, Andreas M. Ernst, Andrew D. Chisholm

## Abstract

The apical extracellular matrix (aECM) of external epithelia often contains lipid-rich outer layers that contribute to permeability barrier function. The external aECM of nematode is known as the cuticle and contains an external lipid-rich layer, the epicuticle. Epicuticlins are a family of tandem repeat proteins originally identified as components of the insoluble fraction of the cuticular aECM and thought to localize in or near epicuticle. However, there has been little *in vivo* analysis of epicuticlins. Here, we report the localization analysis of the three *C. elegans* epicuticlins (EPIC proteins) using fluorescent protein knock-ins to visualize endogenously expressed proteins, and further examine their *in vivo* function using genetic null mutants. By TIRF microscopy, we find that EPIC-1 and EPIC-2 localize to the surface of the cuticle in larval and adult stages in close proximity to the outer lipid layer. EPIC-1 and EPIC-2 also localize to interfacial cuticles and adult-specific cuticle struts. EPIC-3 expression is restricted to the stress-induced dauer stage, where it localizes to interfacial aECM in the buccal cavity. Strikingly, skin wounding in the adult induces *epic-3* expression, and EPIC-3::mNG localizes to wound scars. Null mutants lacking one, two, or all three EPIC proteins display reduced survival after skin wounding yet are viable with low penetrance defects in epidermal morphogenesis. Our results suggest EPIC proteins define specific aECM compartments and have roles in wound repair.

**Highlights:**

- *C. elegans* epicuticlin (EPIC) proteins localize to specific regions in cortical and interfacial cuticle
- Epicuticlins colocalize with BLI collagens in struts in adult cuticle
- EPIC-3 is normally expressed in dauer stage and upregulated by skin wounding
- Mutants lacking all three epicuticlins are viable and show reduced survival after skin wounding

## Introduction

Animal barrier epithelia contain a complex apical extracellular matrix (aECM) that provides structural integrity and forms part of the permeability barrier. Extracellular lipid layers are key components of the matrix permeability barrier, for example the lamellar lipids and cornified lipid envelope of the mammalian stratum corneum (Jonca and Simon, 2023), the tear film lipid layer of the cornea (Pflugfelder and Stern, 2020), or the surfactant lipid layers of alveolar lung cells (Olmeda et al., 2017). The outer lipid layer of arthropods is known as the envelope (Locke, 2001), whereas the equivalent structure in nematodes is termed epicuticle (Bird and Bird, 1991). Extracellular lipid layers can be generated by secretion of flattened lipid disks from organelles such as lamellar bodies (Menon et al., 2018) or from exosome-like vesicles (Tsarouhas et al., 2023). In the vertebrate lung alveolar epithelium lipid-binding proteins such as saposins play key roles in biogenesis of new lipid disks (Sever et al., 2021). However the composition and biogenesis of extracellular lipid layers and their interaction with other aECM compartments in general remain poorly understood.

We are interested in the nematode epicuticle as a model extracellular lipid layer. Most nematode species examined exhibit an epicuticle with smooth trilaminar appearance in TEM approximately 10-30 nm thick. More complex multilaminate or folded epicuticles have been seen in some parasitic species or stages (Franz et al., 1987; Sayers et al., 1984). In other nematodes epicuticle and its lipids have been implicated in host-pathogen interactions (Brivio and Mastore, 2020) and in resistance to abiotic stresses such as desiccation (Bird and Buttrose, 1974; Wharton and Lemmon, 1998). Based on its trilaminar appearance in TEM and the presence of fracture planes in freeze-fracture EM (Peixoto and De Souza, 1995), a prevailing model is that epicuticle resembles a lipid bilayer with as-yet unknown integral proteins.

Imaging, biochemical, and genetic studies confirm that the epicuticle contains lipids. The *C. elegans* epicuticle can be stained by lipophilic dyes (Schultz and Gumienny, 2012) as in other nematodes (Kennedy et al., 1987). Lipophilic dye uptake by the epicuticle is selective and dye mobility is generally low; in parasitic nematodes surface lipid behavior differs between pre-parasitic and parasitic forms (Proudfoot et al., 1993) suggesting epicuticle may be developmentally modulated. Biochemical studies indicate the *C. elegans* epicuticle contains a complex mixture of polar lipids, phosphoglycerides, ceramides, sphingomyelin, and cardiolipin (Bada Juarez et al., 2019; Blaxter, 1993). Moreover, several *C. elegans* lipid biosynthesis mutants display cuticle permeability defects, consistent with epicuticle lipids being required for permeability barrier function (Kage-Nakadai et al., 2010; Loer et al., 2015; Njume et al., 2022; Watts et al., 2003). It remains unclear how the epicuticle lipid layer assembles or how it is attached to the rest of the cuticle.

In biochemical studies of the *C. elegans* cuticle, the epicuticle and underlying outer cortical layer form part of the BME-insoluble and collagenase-resistant fraction (Cox et al., 1981a), suggesting the epicuticle and associated proteins are not collagens. The insoluble fraction has a highly biased amino acid composition both in *C. elegans* (Cox et al., 1981a) and *Ascaris* (Fujimoto and Kanaya, 1973). Likely components of the insoluble fraction include zona pellucida (ZP) family cuticlins or cuticlin-like (CUT, CUTL) proteins (Lassandro et al., 1994; Ristoratore et al., 1994; Sebastiano et al., 1991). Monoclonal antibodies raised against the insoluble fraction of *Ascaris* cuticle were used to identify a novel protein AsCut, later renamed epicuticlin-1, which localized to the epicuticle layer in *Ascaris* and *Brugia* (Bisoffi et al., 1996). *Ascaris* epicuticlin-1 is made up of seven near-perfect Ala and Gly-rich tandem repeats of 49-51 aa, containing two motifs (YGDE and GYR) also found in some insect cuticular proteins (Cornman, 2010).

Proteins related to epicuticlins are found widely in nematodes (Betschart et al., 2022) but their *in vivo* roles remain little understood. Here we analyzed the expression and function of the three *C. elegans* epicuticlin (EPIC) proteins and their relationship to the epicuticle. We document *C. elegans* epicuticle morphology in the wild type using modern EM methods. Using fluorescent protein knockins to endogenous *epic* genes and TIRF microscopy, we show the *C. elegans* epicuticlins are localized to specific aECM substructures, consistent with their localization to epicuticle or to the external cortical layer. Mutants lacking all three epicuticlins are viable with largely normal morphology and permeability barrier function. EPIC-3 expression is normally confined to dauer larvae but is induced in adults by skin wounding and localizes to the wound scar. Moreover, *epic* triple null mutants are defective in adult wound repair. Our results suggest the epicuticlins are not essential for epicuticle biogenesis but may play roles in specific epicuticle regions or in barrier repair.

## Results

### Morphology of the *C. elegans* epicuticle

The *C. elegans* epicuticle has been observed in classic ultrastructural studies (Costa et al., 1997; Cox et al., 1981b), however details of its morphology have not been described at high resolution. We examined the epicuticle in EM of wild type adults and larval stages, using high pressure freeze fixation (HPF) with OsO_4_ to highlight lipid membranes (see Methods for details of samples). The trilaminate epicuticle is defined as two osmiophilic layers separated by an electron-lucent layer. This morphology was consistent in all stages examined (Figure 1A-D). In some specimens the outer osmiophilic layer was less stained than the inner layer; in other samples the outer layer displayed gaps or discontinuities (e.g. Figure 1B). Cortical cuticle staining was often non-uniform, with the outer cortical layer underlying the epicuticle being more electron-dense than the inner cortical layer (Figure 1C).

**Figure 1.**
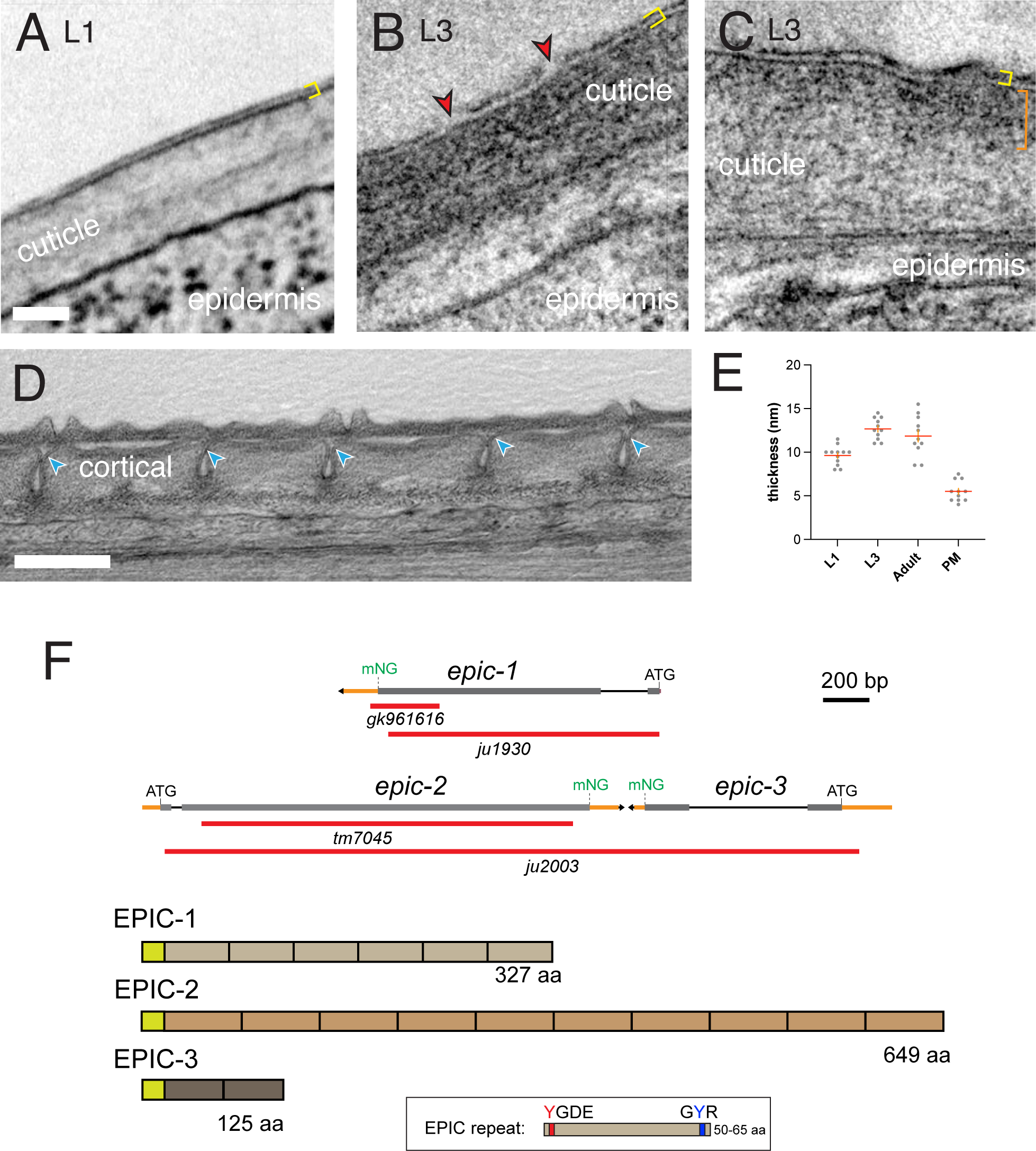
Ultrastructure of the *C. elegans* epicuticle and overview of *C. elegans* epicuticlin (*epic*) genes and proteins. A. Transverse section of L1 cuticle showing L1 epicuticle (yellow brackets). Series SEM_L1_4, z167. Image is 500 nm square; L1 cuticle is ∼150 nm thick. B. Transverse section of L3 cuticle, showing L3 epicuticle, yellow brackets. Series TEM_L3, z202. Image 500 nm square. Red arrowheads indicate discontinuities in the outer osmiophilic layer. C. Example of differential staining of outer cortical cuticle (orange bracket), L3 stage. Images 500 nm square. D. Late L4 stage animal with nascent adult cuticle under detached L4 cuticle, longitudinal section (Adams et al., 2023). Blue arrowheads indicate nascent epicuticle at sides of furrows. Scale, 500 nm. E. Quantitation of epicuticle thickness in EM data, defined as peak-to-peak distance in line scans across the outer osmiophilic layers. PM, plasma membrane thickness as measured in the same sections. F. *epic* gene structures, showing locations of deletions and knock-in insertions. Cartoons of EPIC proteins and the epicuticlin repeat.

In late L4 stages the adult cuticle is synthesized underneath the L4 cuticle. In TEM of late L4 the adult epicuticle is most clearly seen at the sides of nascent furrows and not at the surface of annuli (Figure 1D). From this limited survey epicuticle biogenesis may begin at furrows and later extend to cover annuli, reminiscent of observations of the embryonic epicuticle (then termed the external cortical layer) (Costa et al., 1997). These observations suggest epicuticle forms early in cuticle biogenesis.

We measured epicuticle thickness as the distance between peak electron densities of the two osmiophilic layers, after classical methods for estimating plasma membrane thickness (Yamamoto, 1963). In L1 animals the epicuticle was 9.6 ± 0.3 nm thick (Figure 1A; mean ± SEM, n = 12), whereas later larval stages or adult epicuticle were slightly thicker (10-12 nm, n = 11-14 per stage; Figure 1E). We measured plasma membrane thickness in nearby tissues in these sections to be 5.5 ± 0.3 nm. Based on these measurements the *C. elegans* epicuticle across stages is approximately twice as thick as a typical plasma membrane.

### *C. elegans* EPIC proteins localize to specific compartments of the aECM

*C. elegans* encodes three epicuticlins, defined as secreted proteins composed of two or more 50-65 aa repeats related to those of Ascaris epicuticlin-1 (Figure 1F). Genome-wide transcriptomic studies and gene-specific reporters indicate that *epic-1* and *epic-2* transcripts oscillate strongly in larval development (Meeuse et al., 2020; Meeuse et al., 2023). In contrast *epic-3* transcripts are restricted to dauer or predauer larvae (Contrino et al., 2012). Targeted DamID (TaDa) studies are consistent with *epic-1* and *epic-2* expression in the syncytial epidermis hyp7 and in seam cells in larval stages (Katsanos et al., 2021). *epic-1* and *epic-2* are both enriched in interfacial epidermal cells in adult-specific single cell RNAseq (Ghaddar et al., 2023). Taken together these observations suggest *epic* genes are expressed in multiple epidermal cell types; the oscillating expression of *epic-1* and *epic-2* during larval development is consistent with a role in cuticle synthesis.

We tagged the EPIC proteins by CRISPR/Cas9 mediated insertion of mNeonGreen (mNG) to their endogenous loci at their C-termini (see Methods). EPIC-1::mNG and EPIC-2::mNG were visible in late embryonic stages through to adult stage, becoming brightest in adults. EPIC-1::mNG expression was detectable in the rectal cuticle in late threefold stage embryos (Figure 2A); EPIC-2::mNG was detected in the nose cuticle and faintly in the extraembryonic space. In early L1 larvae EPIC-1::mNG and EPIC-2::mNG were visible in the nose tip and rectal cuticle (Figure 2A, B). From mid-L1 stage onwards EPIC-1::mNG expression was seen in the body cuticle, including L1 alae, consistent with transcriptional reporters showing *epic-1* is expressed at low levels in early L1 then upregulated ∼12 h post hatching (Meeuse et al., 2020). Both EPIC-1::mNG and EPIC-2::mNG were concentrated at the outer ridges of the L1 alae. EPIC-1::mNG localized to cuticle overlying the lateral seam in L2, L3 and L4 stages. In larval body cuticles EPIC-1::mNG and EPIC-2::mNG was most prominent in annuli and excluded from furrows (Figure 2C,D). Within annuli EPIC-1::mNG and EPIC-2::mNG were not uniform but had a granular appearance; small patches or holes lacking EPIC-1::mNG were visible in the mid-lateral cuticle, and in annuli, often aligned with furrows (Figure 2E). In contrast to their oscillating transcription, the fluorescence levels of EPIC-1::mNG and EPIC-2::mNG did not vary strongly within a larval stage, consistent with the proteins being stably incorporated into cuticle. In adults EPIC-1::mNG and EPIC-2::mNG were diffusely localized to annuli and alae; as described below, EPIC-1::mNG and EPIC-2::mNG also became associated with struts in adults.

**Figure 2.**
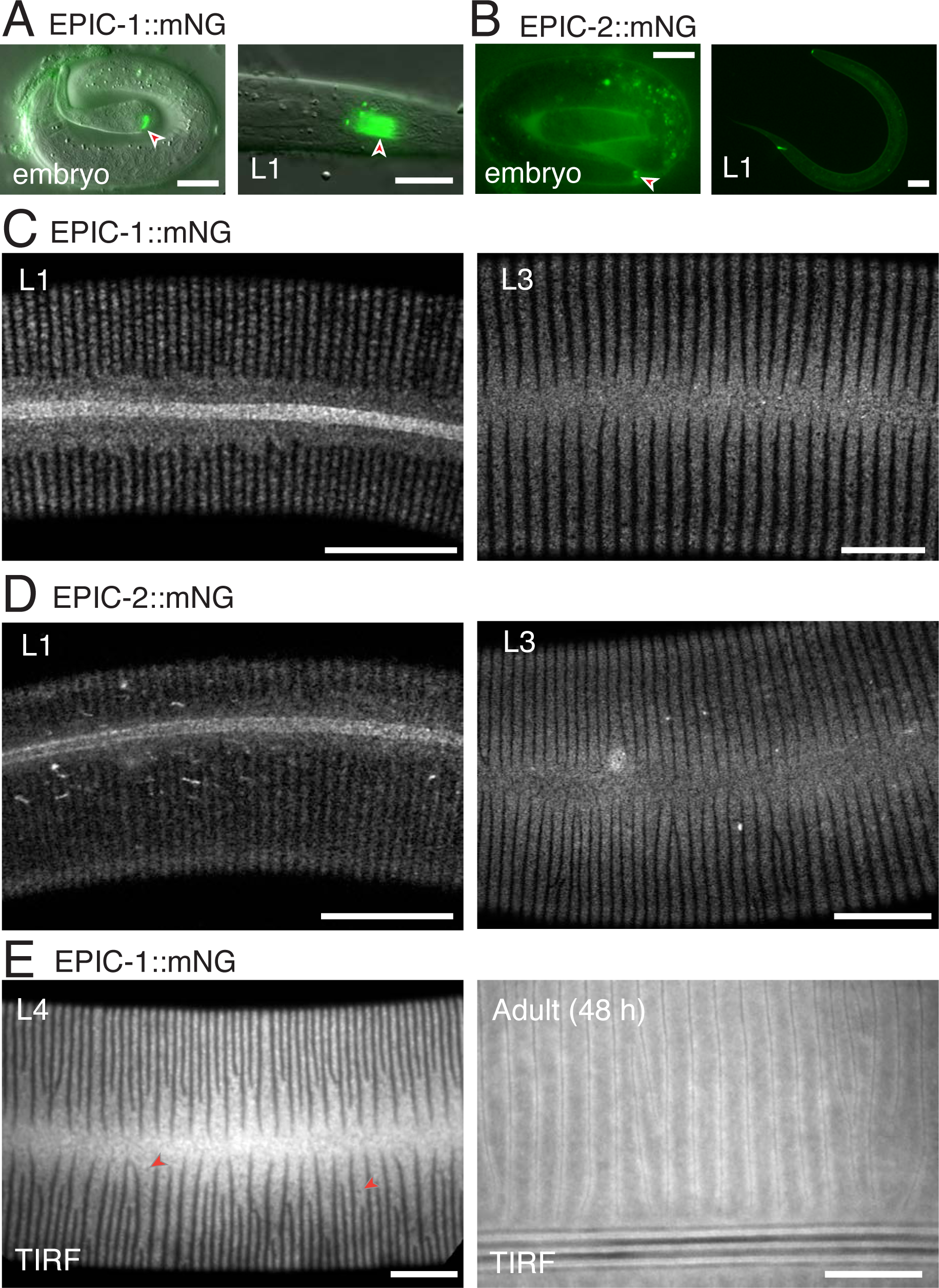
Localization of EPIC-1 and EPIC-2 knock-ins in embryonic, larval and adult aECM A. EPIC-1::mNG expression patterns in embryos and L1s; widefield DIC and fluorescence. EPIC-1::mNG was seen in rectal cuticle and faintly in body cuticle; late threefold embryos. Right, EPIC-1::mNG in L1 larva. EPIC-1::mNG was most intense in distal rectal cuticle, and in two bright dots near the proximal rectum, possibly rectal epithelial cells. Scale, 10 μm. B. EPIC-2::mNG was detected in nose cuticle and faintly in the extraembryonic space of late threefold embryos. In L1 animals EPIC-2::mNG expression was most intense in nose and rectal cuticle and faintly in body cuticle. Scale, 10 μm. C. EPIC-1::mNG in lateral cuticle of L1 and L3 stages, showing localization to L1 alae and granular localization in annuli. Airyscan imaging, scale 10 μm. D. EPIC-2::mNG in lateral cuticle of L1 and L3 stages, showing localization to L1 alae and granular localization in annuli. In larvae and adults EPIC-2::mNG (and to a lesser extent EPIC-1) also localized to puncta or elongated vesicles in deeper focal planes corresponding to transient localization in the secretory pathway of underlying lateral epidermis (not shown). E. EPIC-1::mNG in TIRF microscopy of L4 and adults (48 h post L4). Arrowheads indicate holes in EPIC-1::mNG distribution. Imaging using ring TIRF, surface plane with 100 nm penetration depth. Scale, 10 μm.

To assess whether EPIC proteins localized close to the cuticle surface we tested whether they could be visualized using total internal reflection (TIRF) microscopy. Under stringent TIRF conditions (100 nm penetration depth; see Methods) we were able to visualize diffusely localized EPIC-1::mNG both in L4 stage and in adults (L4+48 h) (Figure 2E; Supplemental Figure 1). As the cuticle in adults is ∼ 1 μm thick, these observations are consistent with EPIC-1::mNG protein localizing near the cortical surface.

In dauer larvae generated by starvation EPIC-1::mNG, EPIC-2::mNG, and EPIC-3::mNG each were highly expressed in the mouth and rectal cuticle, as well as in dauer alae and annuli (Figure 3A). Dauer larvae contain a specialized thickened cuticle, the buccal plug, that occludes the mouth starting ∼0.6 μm from the anterior tip of the nose and extending ∼0.7 μm posteriorly (Albert and Riddle, 1983) (Figure 3B). EPIC-2::mNG localized most anteriorly and could be resolved into longitudinal bars and a thin transverse sheet at the posterior (Figure 3B). These dimensions were consistent with EPIC-2::mNG localization. EPIC-1::mNG localization in dauers resembled that of EPIC-2::mNG (not shown). In contrast, EPIC-3::mNG localized to flat sheets with threefold symmetry that began ∼1.5 μm into the mouth and extended ∼1.5 μm posteriorly (Figure 3B). EPIC-3::mNG may define a distinct compartment in the dauer buccal aECM.

**Figure 3.**
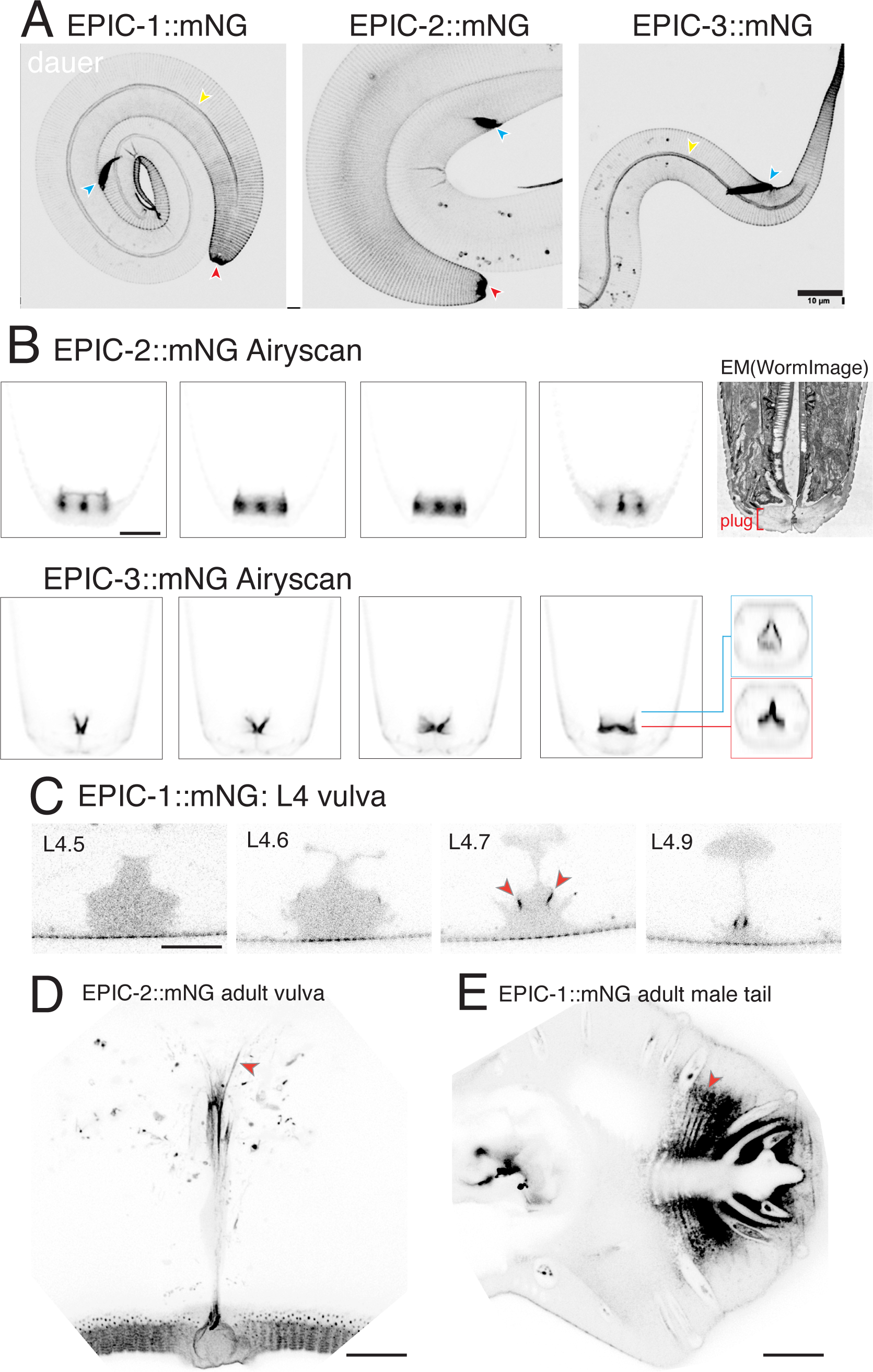
EPIC::mNG knock-in localization in dauer larvae, hermaphrodite vulva, and male tail A. Images of dauer larvae bodies and head region, showing buccal area (red arrowhead) and rectal cuticle (blue arrowheads). EPIC proteins also localized to dauer alae (yellow arrowheads) and to annuli of the body cuticle. Scale 10 μm. B. EPIC-2::mNG and EPIC-3::mNG in dauer buccal region. Single focal planes of Airyscan z stacks, panels 10 x 10 μm, Scale 2 μm. EPIC-2::mNG localizes to anterior mouth, EPIC-3::mNG localizes to posterior mouth. Orthogonal sections of EPIC-3::mNG are 1 μm apart (indicated by colored lines). The TEM image of the dauer buccal plug is from WormImage N2_dauer_50-7-2_34. C. EPIC-1::mNG in L4 morphological substages; diffuse localization in vulval lumen and near the surface of vulC cells by L4.7 (blue arrowheads). 30 x 20 μm, single focal plane, LSM800 confocal. D. EPIC-2::mNG localizes to filamentous structures (red arrowheads) in the adult vulva, projection of 5 deep focal planes, LSM800 confocal. Strain is in *epic-1(0)* background. Scale, 10 μm. E. EPIC-1::mNG in adult male tail, ventral view. EPIC-1::mNG is diffuse in the male tail fan and enriched in ribbed structures (red arrowhead) in the posterior fan, similar to BLI-1::mNG (Adams et al., 2023). EPIC-1::mNG signal is also visible inside sensory rays. Scale, 10 μm.

As well as being localized to the main body cuticle generated by hyp7 and seam cells, EPIC-1::mNG and EPIC-2::mNG were strongly expressed in cuticle generated by interfacial epidermal cells (e.g. the nose tip, rectum, vulva). In L4 substage 4.5-4.9 animals (Mok et al., 2015), EPIC-1::mNG localized diffusely in the vulval lumen as well as on the luminal surface near vulC cells, indicating that it was secreted from the vulval epithelium and localized to the developing vulval cuticle (arrowheads, Figure 3C). In adults EPIC-1::mNG became more widely localized to the apical surface of the vulva and was also observed in filamentous structures within the vulva (Figure 3D). EPIC-1::mNG localization in the male body cuticle was similar to that in hermaphrodites; in the adult male tail EPIC-1::mNG and EPIC-2::mNG were diffusely localized in the cuticular fan, most intensely in the posterior fan within which they had a ribbed appearance (arrowhead, Figure 3E).

To understand the biochemical properties of EPIC::mNG proteins, we made cuticle preparations and obtained soluble fractions. By Western blot analysis, we detected mNG-tagged proteins for both EPIC-1::mNG and EPIC-2::mNG in preparations of soluble cuticle proteins from mixed stages (Supplemental Figure 2A). Faint bands could be seen at sizes that may correspond to monomeric proteins, however most signal was observed in high-molecular weight species or smears of ∼150 kDa and larger. As EPIC proteins and their mNG fusion proteins are highly acidic (e.g. EPIC-1::mNG has a pI 4.46) they may migrate slower than predicted in SDS-PAGE. These observations validate the expression of mNG-tagged EPIC proteins as components of cuticle.

### Loss of function in *epic* genes has mild effects on cuticle morphology and function

We generated *epic* null mutations by CRISPR/Cas9 mediated gene editing and examined additional deletion alleles generated by knockout projects (Table 1). *epic-1(gk961616)* is a 307 bp deletion in the 3’ end of *epic-1*, eliminating half of repeat 5 and all of repeat 6 as well as 34 bp of the 3’ UTR. We assayed *epic-1* transcript structure by RT-PCR (Supplemental Figure 2B). However, due to the repetitive sequence nature of *epic-1* and *epic-2*, RT-PCR of *epic-1* or *epic-2* generated multiple bands in wild type animals. *epic-1(gk961616)* mutants expressed a deleted transcript that could encode a truncated protein. We therefore generated a large deletion mutation of *epic-1* using CRISPR/Cas9 (see Methods). *epic-1(ju1930*) null mutants were viable and fertile with normal brood size. *epic-2(tm7045)* is predicted to delete repeats 2-10 and cause a frameshift in repeat 1; we verified that *tm7045* mutants expressed truncated transcripts (Supplemental Figure 2B). Mutations in *epic-3* were not available. *epic-2* and *epic-3* genes are immediately adjacent in tail-to-tail orientation (Figure 1F). As their repeated sequences limited options for crRNAs we generated a deletion eliminating most of both genes, *epic-2&3(ju2003)*. These mutants were viable with low penetrance lethality (Figure 4A), fertile, and had wild type body shape.

**Table 1.** Molecular description of *epic* mutants and compound mutants.

| Genotype | Genomic DNA | Predicted protein | Brood size (mean ± SEM) |
| --- | --- | --- | --- |
| *epic-1(gk961616)* | 307 bp deletion | Truncated in repeat 5 after V255 | ND |
| *epic-1(ju1930)* | 1187 bp deletion | Deletes entire coding sequence. | 173 ± 46 |
| *epic-2(tm7045)* | 1776 bp deletion | Truncated after repeat 1 | 176 ± 48 |
| *epic-2&3(ju2003)* | 3335 bp deletion | Deleted all of *epic-2* and *epic-3* coding sequence. | 210 ± 63 |
| *epic-1(gk961616) epic-2(tm7045)* | | | 37 ± 20 |
| *epic-1(ju1930) epic-2(tm7045)* | | | 162 ± 49 |
| *epic-1(ju1930) epic-2&3(ju2003)* | | | 170 ± 35 |

n = 10 broods counted per genotype.

**Figure 4.**
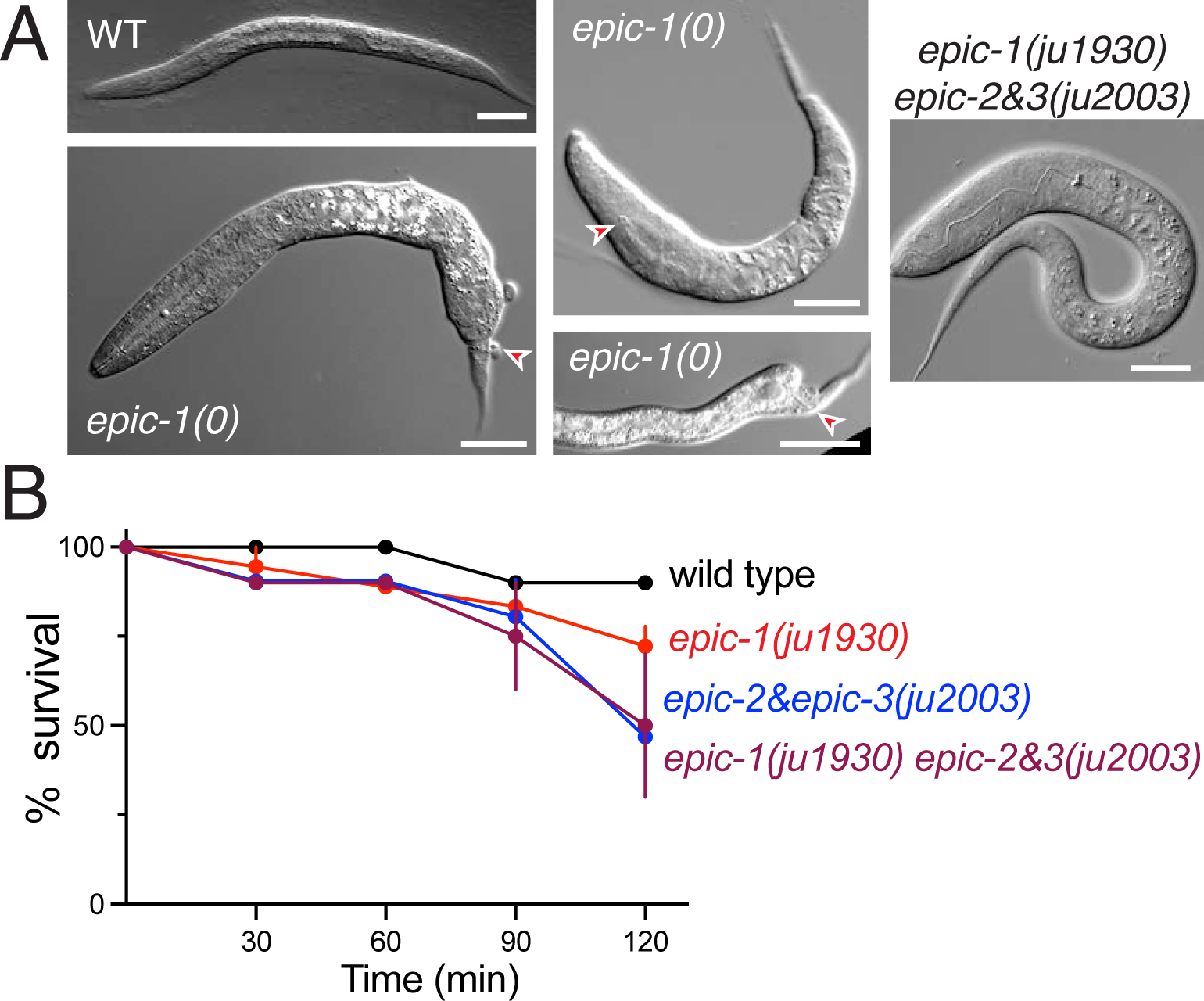
*epic* loss of function mutant phenotypes A. L1 morphology in wild type and examples of low penetrance phenotypes observed in *epic-1(ju1930)* and in *epic-1(ju1930) epic-2&3(ju2003)* mutants. Examples (red arrowheads) of tail Vab, unattached pharynx (Pun) and blocked rectum from *ju1930*. Example of L1 arrested *epic-1(ju1930) epic-2&3(ju2003)* animal. Scales, 20 μm. B. Cuticle permeability in wild type and *epic* mutants, desiccation assay. n = 4-6 sets of 10 animals per time point.

To investigate potential functional redundancy between *epic-1* and *epic-2* or *epic-3* we generated double and triple mutants by recombination. *epic-1(ju1930) epic-2&3(ju2003)* compound mutants were viable and fertile with normal cuticle morphology (e.g. alae, annuli, and furrows as assessed by DIC) and low penetrance epidermal morphology defects such as lumpy body shape or unattached pharynxes (Figure 4A). *epic-1(gk961616) epic-2(tm7045)* double mutants displayed reduced fertility (Table 1) and overall slow growth to adults. The phenotypes observed in *epic-1(gk961616) epic-2(tm7045)* strains may be due to unidentified background mutations as they were not observed in the triple null mutant. Alternatively these defects may be due to the expression of truncated EPIC-1 proteins made in the *epic-1(gk961616)* mutant.

We used RT-PCR to examine whether *epic* transcript levels displayed compensation (Supplemental Figure 2B). *epic-1* transcripts were expressed at normal levels in *epic-2(tm7045)* and in *epic-2&3(ju2003)*. Conversely *epic-2* transcripts appeared normal in *epic-1(ju1930)*, although it was not possible to establish a quantitative difference due to variable priming from internal repeats. Based on these data *epic* transcripts do not display significant compensation.

We addressed whether EPIC proteins affected each other’s localization. EPIC-1::mNG localization in *epic-2(tm7045)* L4 stage appeared indistinguishable from normal (not shown); conversely EPIC-2::mNG localization in *epic-1(gk961616)* resembled that of the wild type (Supplemental Figure 2C). Together these findings suggest EPIC-1 and EPIC-2 do not regulate each other’s localization.

Given their localization to epicuticle, we assessed cuticle permeability barrier function in *epic* mutants and observed mild cuticle permeability defects compared to barrier function mutants such as *gmap-1* (Njume et al., 2022). For example in assays of sensitivity to desiccation 63% of *epic-1(ju1930) epic-2&3(ju2003)* were inviable after 60 min compared to 68% of wild type or 0% of *gmap-1(0)* mutants (n = 40-60 per genotype, Figure 4B). Overall these results suggest loss of EPIC function has minimal effects on the cuticle permeability barrier.

### EPIC-1 and EPIC-2 localize to adult struts, dependent on BLI collagens

Struts are adult-specific columnar structures that connect cortical and basal cuticle layers, spanning the fluid-filled medial layer (Adams et al., 2023). Loss of function in strut collagens leads to separation of cuticle layers (‘blistering’). Although *epic* mutants did not display overt blistering, EPIC-1::mNG and EPIC-2::mNG both displayed adult-specific puncta resembling strut puncta as defined by the BLI collagens (Figure 5A). The punctate localization of EPIC-1::mNG and EPIC-2::mNG was in addition to their diffuse localization in cortical cuticle; in confocal z series EPIC-1::mNG or EPIC-2::mNG puncta were much brighter than the diffuse cortical fluorescence seen in younger adults, suggesting they may reflect new EPIC protein synthesis in the adult.

**Figure 5.**
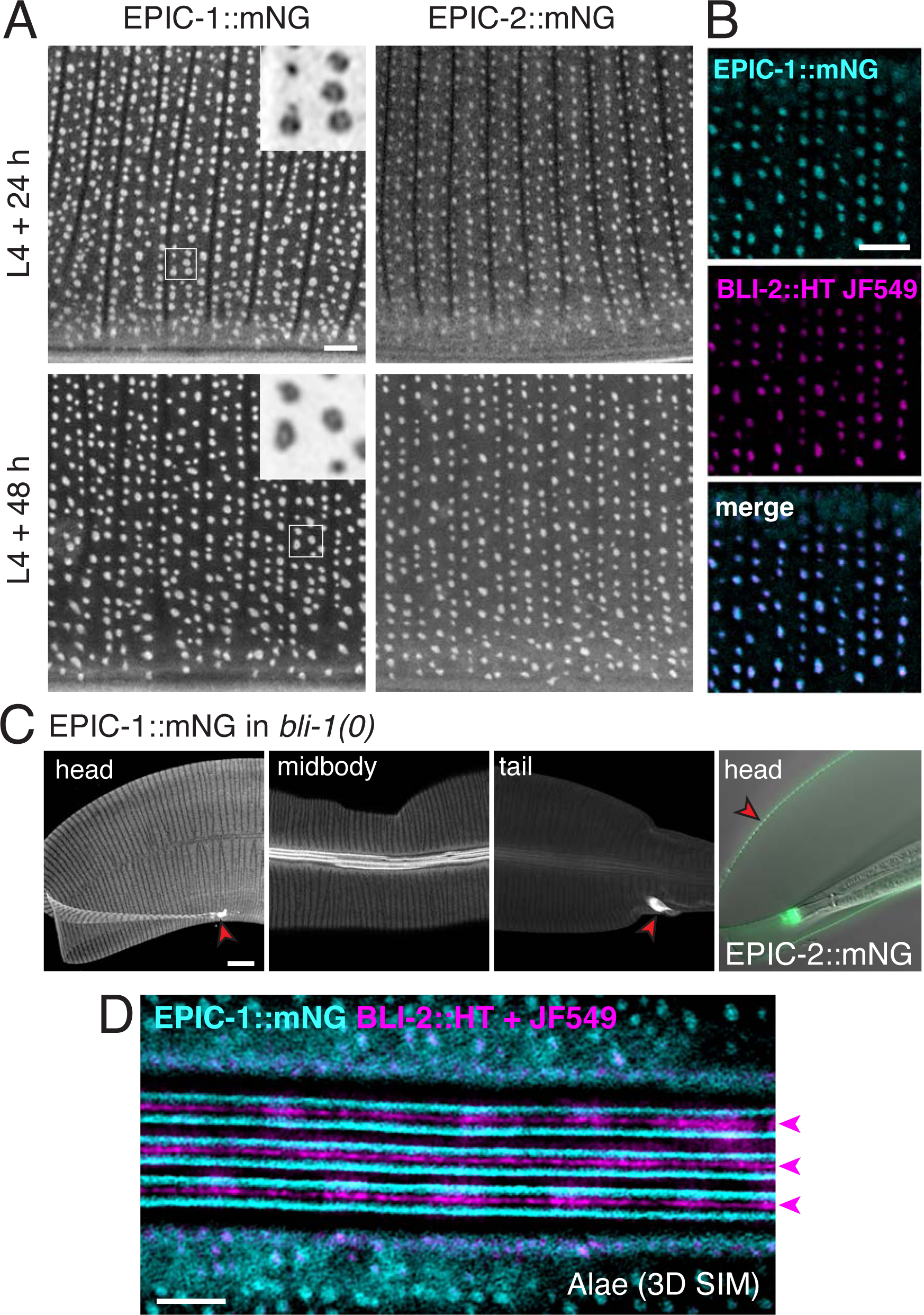
Localization of EPIC proteins to adult struts and alae A. EPIC-1::mNG and EPIC-2::mNG in adults 24 h and 48 h post L4, anterior body lateral epidermis, Airyscan. Main images are 20 x 20 μm. Insets showing donut morphology are 2 x 2 μm. Scale, 2 μm. B. Colocalization of EPIC-1::mNG (cyan) and BLI-2::HT stained with JF549 dye (magenta) in 3D SIM, single focal plane. Panels 10 x 10 μm. Scale, 2 μm. C. EPIC-1::mNG in *bli-1(ju1395)* null mutant adult: head, midbody, and tail regions. EPIC-1::mNG localizes to annuli, alae, and interfacial cuticle of nose and rectum (red arrowheads) but not to struts. Images 70 x 70 μm. Right panel, EPIC-2::mNG localization in *bli-1(ju1395)*, DIC and fluorescence showing localization to outer surface of blistered cuticle (red arrowhead) and to buccal cuticle. Scale, 10 μm. D. BLI-2::HT stained with JF549 dye (magenta) localizes to adult alae ridges (magenta arrowheads) and EPIC-1::mNG (cyan) and localizes to the sides of ridges. Maximum intensity projection of 3D SIM reconstruction, image is 20 x 10 μm. Scale, 2 μm.

Struts form in the late L4 stage. Although EPIC::mNG proteins were expressed through L4.5-L4.9 they remained diffuse in the cortical cuticle and were not localized to struts (Supplemental Figure 3A); faint localization to longitudinal alae could be seen by L4.8. By examining staged adults we found that EPIC-1::mNG and EPIC-2::mNG became recruited to struts beginning ∼12 h after the L4/Adult molt. These observations indicate the EPIC proteins are not localized to struts at the time of initial strut formation but are recruited to struts in early adult life. EPIC-1::mNG and EPIC-2::mNG could also be visualized in the epidermal secretory pathway (i.e. puncta or filaments in deeper focal planes; not shown) until at least 48 h in adulthood, suggesting EPIC proteins are secreted by the epidermis through early adulthood.

We focused on EPIC-1::mNG for quantitative analysis of strut localization because EPIC-1::mNG displayed a sharp transition from diffuse to punctate compared to EPIC-2::mNG. The patterning of EPIC-1::mNG puncta resembled those of BLI-1 puncta, forming three circumferential rows per annulus (two furrow-flanking rows and a more variable central row) (Figure 5A). The spacing and density of EPIC-1::mNG puncta also resembled that of BLI-1. For example in furrow-flanking rows EPIC-1::mNG puncta were an average of 0.81 ± 0.02 μm apart (mean ± SEM, n =10 rows). compared to 0.77 μm for BLI-1::mNG. The density of EPIC-1::puncta in midbody lateral cuticle was 153 puncta per 100 μm^2^ (n = 6 ROIs) compared to 145-183 puncta per 100 μm^2^ for BLI-1::mNG.

We previously showed that struts contain the three BLI collagens BLI-1, BLI-2, and BLI-6. We found that EPIC-1::mNG and BLI-2::HaloTag (HT) colocalized in adult struts (Figure 5B). EPIC-1::mNG and BLI-1::mScarlet (mSc) also displayed significant colocalization in adults but not in L4 (Supplemental Figure 3B). EPIC-1::mNG and BLI-1::mSc displayed a mean Pearson colocalization coefficient of +0.03 at L4 + 12 h prior to EPIC-1 puncta formation, increasing to +0.52 at L4 + 24 h after EPIC puncta formation (n = 6 ROIs per time point, Supplemental Figure 3C), similar to the degree of colocalization of BLI-1::mSc and BLI-2::mNG (Adams et al., 2023).

Our 3D Structured Illumination Microscopy (SIM) analysis of BLI-1 and BLI-2 knock-ins revealed that both collagens show cylindrical localization in struts, appearing donut-shaped in single z sections (Adams et al., 2023). We performed 3D SIM on EPIC-1::mNG in adult stages and observed donut-shaped structures at struts, however the SIM reconstruction quality was low due to the diffuse localization of EPIC signal in the cortical layer. We therefore estimated EPIC donut size from Airyscan images (Figure 5A). In line scans of EPIC-1::mNG puncta displaying donut morphology, meaning a central minimum surrounded by peaks, EPIC-1::mNG peak to peak diameter was 239 ± 47 nm in adults 24 h post L4 (mean ± SEM, n = 11) and 280 ± 63 nm in adults 48 h post L4 (n = 11) compared to our SIM measurements of BLI-1::mNG peak to peak diameter of 160 nm. These observations suggest the EPIC donuts may be slightly larger in diameter than the BLI donuts, consistent with EPIC proteins being recruited to the outer layer of struts.

We next addressed whether EPIC::mNG localization in struts depended on *bli* collagen. In *bli-1(ju1395)* null mutants which lack struts and have an expanded medial layer (‘blister’) (Adams et al., 2023), EPIC-1::mNG formed a diffuse granular pattern in annular bands at the outer surface of the blister and did not form strut-like puncta at any stage (Figure 5C); EPIC-1::mNG localization to alae or interfacial cuticle was not affected. EPIC-2::mNG was also localized to the surface of blistered cuticle in *bli-1(0)* (Figure 5C) and occasionally accumulated diffusely within the fluid blister. Together these observations suggest EPIC proteins require struts for their punctate localization, and that in the absence of struts they remain diffuse in the cortical layer of the cuticle. Conversely, BLI-1 or BLI-2 localization in *epic-1(0)* or *epic-2(0)* mutants was indistinguishable from WT (not shown), consistent with the lack of Bli phenotypes in *epic* mutants.

Although the diffuse localization of EPIC-1::mNG precluded high quality 3D SIM reconstructions of struts, we were able to visualize the localization of EPIC-1::mNG and BLI-2::HT at adult alae. BLI-2::HT formed three longitudinal stripes corresponding to the ridges of the alae, whereas EPIC-1::mNG could be resolved into six longitudinal stripes flanking the BLI-2-expressing ridges (Figure 5D). EPIC-1::mNG also localized to more diffuse longitudinal bands flanking the alae and deeper in the cuticle. These observations indicate that BLI-2 and EPIC-1 localize to distinct parts of the adult alae, with EPIC-1 lining the sides of the alae ridges.

### Cortical EPIC-1::mNG localize close to the epicuticle lipid layer

Our confocal and TIRF imaging indicated EPIC::mNG knock-ins localize to a surface layer of cuticle. To assess their localization relative to the lipid-rich epicuticle we stained EPIC-1::mNG and EPIC-2::mNG animals with the lipophilic dye R18 (Octadecyl Rhodamine B Chloride; see Methods) (Proudfoot et al., 1993). R18 staining, like that of the carbocyanine dye DiI, is intense in furrows, weaker in annuli, and also stains structures in adult alae. In analysis of z sections from stacks with step size 0.29 μm the R18 signal could be detected in 2-3 cortical sections of which 1-2 overlapped with that of diffuse EPIC-1::mNG or EPIC-2::mNG (Figure 6A). As the epicuticle lipid layer is ∼10 nm in depth, it was not possible to resolve in these confocal series whether the EPIC signal is in the same z layer as lipid dye staining or whether it is within an adjacent underlying cortical layer. R18 epicuticle staining in *epic* triple mutant adults was typically less intense than in wild type but was otherwise similar in distribution (Figure 6B).

**Figure 6.**
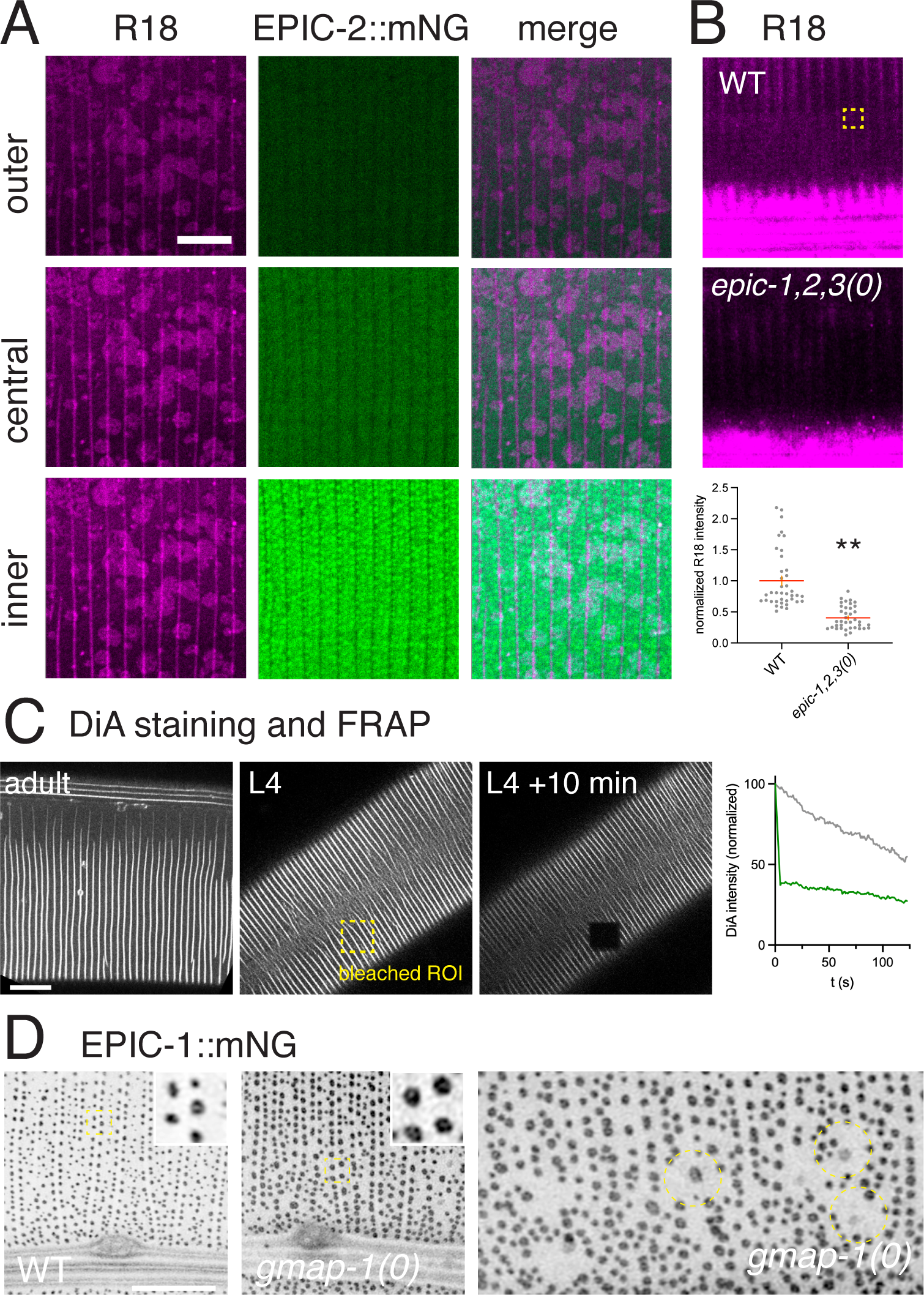
Colocalization of EPIC proteins with lipophilic dyes and interactions with lipid synthesis genes A. R18 staining (magenta) and EPIC-2::mNG (cyan). 3 adjacent z sections Panels are 15 x 15 μm. B. R18 staining in wild type (N2) and *epic-1(ju1930) epic-2&3(ju2003)* triple mutant young adults (L4 + 24 h). Images are maximum intensity projections of 5 focal planes of cuticle staining over lateral epidermis, with the central z plane in the middle of the alae (bottom of images), panels 30 x 30 μm; Scale, 10 μm. B. Quantitation of mean R18 brightness in 2 x 2 μm ROIs of MIPs, P < 0.001 by Mann-Whitney test. C. DiA staining in adult cuticle and DiA FRAP experiment in L4 stage, images pre bleach and 10 min post bleach. Graph shows DiA fluorescence over 2 min post bleach, normalized to 0 = 100%; gray line shows background photobleaching. Panels are 50 x 50 μm. D. EPIC-1::mNG puncta morphology in wild type adult and *gmap-1(ulb13)* null mutants. Anterior epidermis showing anterior deirid sensilla and alae at bottom. Panel 20 x 20 μm, enlarged insets 2 x 2 μm. Right panel, examples of EPIC-1::mNG ‘crop circles’ in *gmap-1(0)* mutants (circular dashed outlines), panel 20 x 10 μm.

Epicuticle lipids were reported to show little lateral mobility in other nematodes based on FRAP experiments (Kennedy et al., 1987). We assessed this in *C. elegans* using staining with the dialkylaminostyryl dye DiA (4-Di-16-ASP) and found that bleached areas in L4 or adult animals showed minimal recovery after 10 min, consistent with low mobility of epicuticle lipids (Figure 6C). EPIC-1::mNG fluorescence did not recover after photobleaching either in larvae or adults (data not shown), suggesting EPIC-1 is stably localized within the cuticle.

The lipid metabolic enzyme alkylglycerol monooxygenase AGMO-1 (Loer et al., 2015) and the GM2AP-like lipid transporter GMAP-1 (Njume et al., 2022) have been implicated in epicuticle lipid biogenesis. Loss of function in *agmo-1* confers mild cuticle permeability defects whereas loss of function in *gmap-1* causes severe permeability defects and pleiotropic defects in cuticle structure. In *gmap-1(ulb13)* null mutants EPIC-1::mNG puncta were normally patterned but larger than in the wild type, such that more puncta had donut morphology. Although the peak-to-peak diameter was not significantly increased (276 ± 32 nm, n = 12), the fraction of an ROI occupied by puncta was significantly increased (24% vs 16.9% in wt, P = 0.0003 by Student’s t test, n = 9 ROIs). Thus, EPIC-1::mNG puncta appeared to be more spread out in *gmap-1(0)* mutants versus wild type*. gmap-1(0)* mutants also displayed scattered ‘crop circles’ containing a larger less intense EPIC-1::mNG punctum surrounded by a region ∼ 2 μm diameter lacking EPIC-1::mNG puncta (Figure 6D). In *agmo-1(0)* null mutants EPIC-1::mNG displayed similar ‘crop circles’ though at lower penetrance (not shown). Taken together these observations suggest EPIC-1::mNG diffuse localization in the cortical cuticle is not significantly affected by loss of *agmo-1* or *gmap-1*; however cuticle lipids may affect strut morphology or EPIC distribution at struts.

### EPIC-3 is upregulated in response to wounding and localizes to the wound scar

Consistent with the lack of EPIC-3 expression in adults, *epic-3* transcripts are undetectable in adults under normal conditions. However, after needle wounding, *epic-3* mRNA was highly upregulated, whereas *epic-1* and *epic-*2 transcript levels were slightly reduced (Fu et al., 2020). We thus examined whether EPIC proteins may be involved in cuticle repair after wounding. EPIC-1::mNG and EPIC-2::mNG showed occasional localization to diffuse rings around the wound site (Figure 7A). In contrast EPIC-3::mNG was upregulated and showed specific localization to ring structures by 6 h post wounding; by 24 h EPIC-3::mNG rings typically had contracted to central puncta, concomitant with wound closure (Figure 7A). As EPIC-3::mNG was not detectable in unwounded adults, this is consistent with *epic-3* transcriptional upregulation after wounding. Animals with mutations in the death associated protein kinase *dapk-1*/DAPK constitutively upregulate multiple wound-induced proteins including antimicrobial peptides and the BLI collagens (Chuang et al., 2016; Tong et al., 2009). EPIC-3::mNG was not detectably upregulated in *dapk-1(ju4)* mutants (not shown), suggesting EPIC-3 regulation is independent of the pathways controlled by DAPK-1.

**Figure 7.**
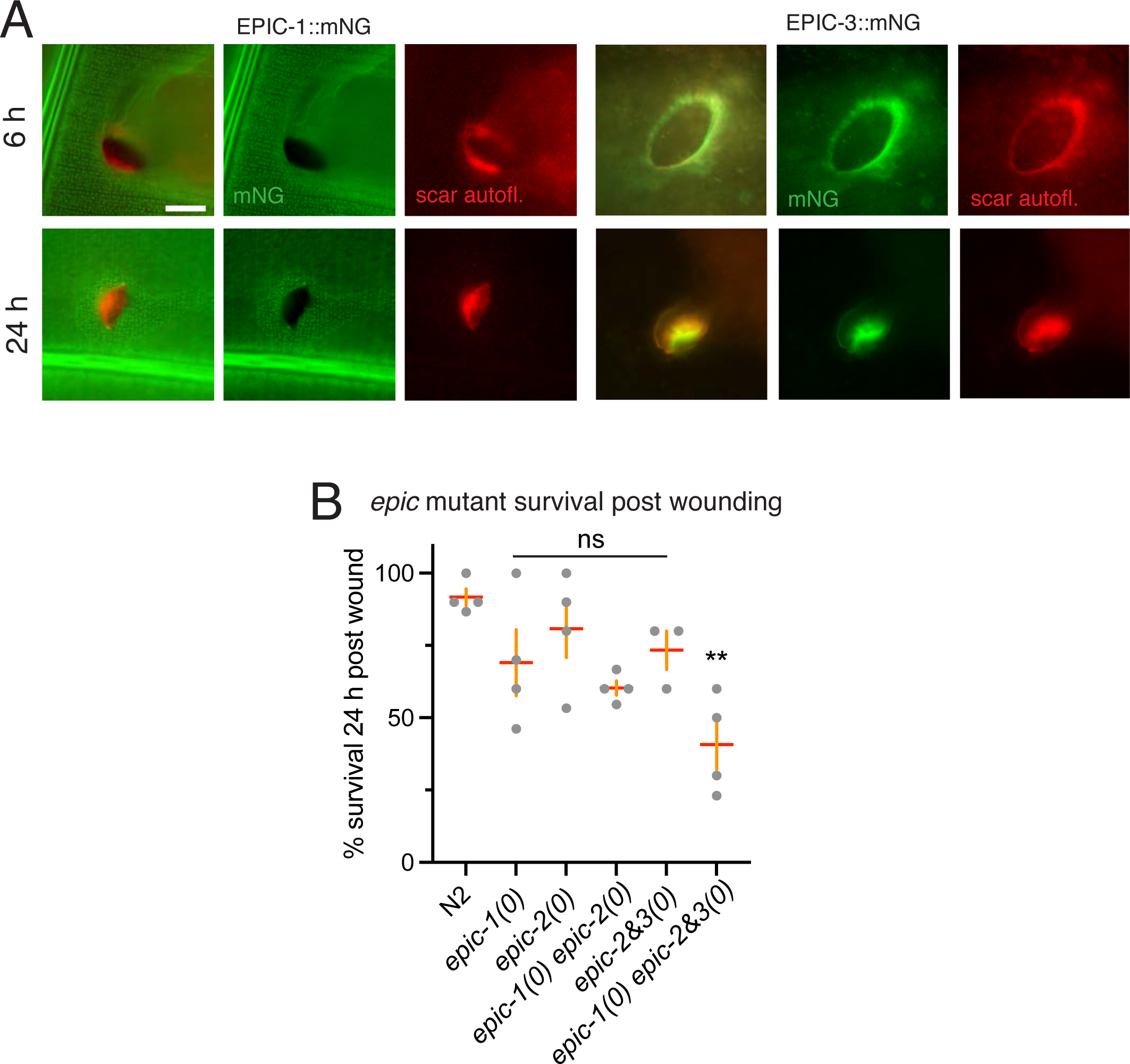
EPIC proteins in wound repair A. EPIC-1::mNG and EPIC-3::mNG localization at wound sites 6 h and 24 h post needle wounding. Needle wounding results in a 5-10 μm diameter hole in the adult EPIC-1::mNG pattern. In later time points EPIC-1::mNG occasionally showed diffuse localization around the wound site but did not colocalize with the endogenous scar autofluorescence (red channel). EPIC-2::mNG after wounding (not shown) resembled EPIC-1::mNG. EPIC-3::mNG localized to closing rings at 6 h and closed puncta at 24 h, typically colocalizing with scar autofluorescence. Widefield microscopy, all panels 40 x 40 μm; Scale, 10 μm. B. Survival of *epic* mutant strains post wounding. Animals were wounded at L4+24 h and checked at L4 + 48h; 100% of animals survived to L4+48 h in the unwounded condition. Single and double mutants showed reduced survival although this was not statistically significant; survival of the *epic* triple mutant was significantly reduced (P = 0.0067 by Kruskal-Wallis test).

To address the function of EPIC proteins in wound repair we examined survival rates of mutants at 24 h after wounding. *epic* single or double mutant survival was reduced although this was not significantly different from that of the wild type, however the *epic-1(0) epic-2&3(0)* triple mutants displayed significantly reduced survival (Figure 7B). Together these results suggest EPIC-3 is recruited to damaged cuticle sites for barrier repair and that EPIC proteins are functionally important for survival after wounding.

## Discussion

Lipid-rich extracellular layers are widely found in the aECM of barrier epithelia from invertebrates to humans, yet their organization and interaction with other aECM constituents remains little understood. Here we focused on the *C. elegans* epicuticlins, which we find localize to specific aECM substructures in the vicinity of the epicuticle. EPIC proteins also localize to other aECM substructures that may have specialized epicuticles. Genetic elimination of epicuticlin function has minimal effects on cuticle morphology, suggesting the epicuticlins may act redundantly or that their absence is compensated for by other factors. EPIC-3 displays most restricted expression, is upregulated by skin wounding, and may specifically play a role in wound repair.

### The *C. elegans* epicuticle and epicuticlins

Our analysis of available HPF fixed EM sections is consistent with classical TEM studies of *C. elegans* that have suggested the epicuticle, previously known as the external cortical layer resembles a thickened lipid bilayer (Cox et al., 1981a). Our measurements of epicuticle thickness are also within the range of thicknesses reported for other nematode epicuticles, which range from 6 to 40 nm (Bird, 1980). Our FRAP imaging of lipophilic dyes supports the model that *C. elegans* epicuticular lipids do not freely diffuse. This lack of lateral mobility has been ascribed to ceramides, although these make up small fraction of epicuticle lipids based on recent lipidomics (Njume et al., 2022).

Epicuticlins were first identified as antigens present in the insoluble fraction of *Ascaris* cuticle (Bisoffi et al., 1996). Our analysis of *C. elegans* EPIC expression is broadly consistent with their localization to the epicuticle and/or the underlying external cortical layer, corresponding to the insoluble fraction of the cuticle. EPIC proteins are made up of low-complexity repeats with a biased amino acid composition. EPIC-1 and EPIC-2 together contain ∼34.5% Alanine, excluding their signal peptides. In contrast, other proteins shown or predicted to be in the insoluble fraction such as CUT proteins are <10% Ala, with the exception of the tandem repeat protein CUT-2 (27% Ala) (Lassandro et al., 1994). EPIC proteins might account for the biased amino acid composition of the insoluble cuticle fraction (19.5% Ala, vs 11% in soluble fraction). The EPIC proteins are also unusually acidic. Highly acidic proteins have been implicated in biomineralized matrices (Marin and Luquet, 2007); as nematode cuticles are not biomineralized, the significance of the highly acidic nature of epicuticlins is not yet clear. The *C. elegans* surface is negatively charged at neutral pH, attributed to sulfolipids or sulfated sugars (Blaxter, 1993). EPIC proteins may potentially contribute to the charged surface of *C. elegans*. Epicuticlins were originally identified via antibodies raised against the insoluble cuticle fraction, yet we could detect EPIC::mNG fusion proteins in extracts of soluble cuticle fractions. It is possible the mNG tag affects the solubility of the EPIC fusion protein; alternatively EPIC proteins may be present in both soluble and insoluble fractions.

We find that EPIC::mNG knock-in proteins localize to an external cortical region of the cuticle close enough to the surface that they can be visualized using TIRF microscopy. Prior TIRF microscopy studies in *C. elegans* have been limited as the cuticle or eggshell are thicker than a TIRF field. Instead, studies have used near-TIRF/semi-TIRF with penetration depths of ∼500 nm (Robin et al., 2014). Our ability to detect cortical EPIC-1::mNG signal under stringent TIRF conditions suggests the outermost EPIC-1 signal is within 100 nm of the outside of the animal, consistent with localization to the epicuticle or external cortical layer.

Our imaging is consistent with EPIC proteins localizing close to the lipid rich layer. In larvae the EPIC-1::mNG and EPIC-2::mNG proteins are localized in annular ridges whereas lipophilic dyes more consistently stain annuli and furrows, suggesting EPIC proteins do not uniformly scaffold a lipid layer. Moreover, in EM studies the epicuticle is first visible within furrows and only later extends to annuli. These observations argue against EPIC proteins being integral parts of the lipid layer. Potentially EPIC proteins may define the external cortical layer in annuli.

### EPIC proteins localize to multiple aECM compartments

EPIC proteins are also highly expressed in interfacial cuticle areas such as nose or rectum. In single cell transcriptomic studies of embryos (Packer et al., 2019) *epic-1* was expressed in interfacial epidermal cells including hyp1, hyp2, and rectal epithelia, consistent with EPIC-1 being secreted by underlying epithelia and being locally incorporated into cuticle. Cuticle overlying interfacial epithelial cells or glial cells has a distinct composition and ultrastructure compared to body cuticle (Fung et al., 2023). The ultrastructure of the epicuticle or its boundaries in interfacial regions have not been extensively characterized in *C. elegans* though an electron-lucent layer is visible in the rectal cuticle (www.wormatlas.org, Slidable Worm, section SW682). In other nematode species the epicuticle is discontinuous and terminates within interfacial regions (Bird, 1980; Dick and Wright, 1974). Speculatively, EPIC proteins may play roles in reinforcing or sealing the epicuticle at certain boundary regions.

Our findings that *epic* single and compound mutants are largely viable and fertile with grossly normal cuticle morphology raises the question of their normal role in development. We have observed low penetrance defects in *epic* mutant epidermal morphogenesis, specifically detachment of the pharynx or blockage of the rectum, suggestive of roles in these interfacial cuticle regions. Notably, several aECM proteins have been implicated in pharynx attachment to the buccal cavity, including fibrillin/FBN-1 (Kelley et al., 2015), DEX-1 (Flatt et al., 2019) and DYF-7 (Heiman and Shaham, 2009). The pharynx-buccal cavity attachment may be especially sensitive to aECM defects.

Interestingly, EPIC-2 and EPIC-3 localize to distinct parts of the buccal cuticle of the dauer larva, with EPIC-2 localizing anterior to EPIC-3. Little is known about the biogenesis or composition of the buccal plug, however it has a highly osmiophilic luminal surface in TEM (Albert and Riddle, 1983) that may be related to the body epicuticle. Our observations suggest EPIC-3 may define a distinct aECM posterior to the buccal plug. As EPIC-3 is also specifically upregulated by wounding and localizes to wound scars, aspects of the dauer-specific aECM program may be reactivated by wounding. EPIC-3 may define a specific aECM compartment that functions to seal orifices either in the dauer mouth or at a wound site.

Unexpectedly, EPIC-1 and EPIC-2 both become localized to struts beginning in adulthood. These observations suggest struts undergo maturation during adult life, with early struts (as generated in L4) being composed of the collagens BLI-1, BLI-2, and BLI-6 and adult struts further recruiting EPIC-1 and EPIC-2. Molecular epistasis indicates BLI proteins are required for EPIC proteins to localize to struts and not the reverse. Moreover, super resolution microscopy reveals that EPIC proteins also display nanoscale organization into cylindrical structures. As our EPIC and BLI knock-ins differ in intrinsic brightness and in their level of diffuse signal, quantitative comparisons in puncta size using different fluorophores should be treated with caution, especially as these comparisons focus on large puncta displaying a central minimum (donut morphology). We note that under similar Airyscan microscopy parameters BLI-1::mNG donuts are not resolvable whereas EPIC-1 donuts are visible in the wild type and even more so in *gmap-1* mutants. Taken together these comparisons suggest EPIC proteins may be recruited to the outside of BLI-containing struts. As *epic* mutants do not display Bli phenotypes, the role of EPIC proteins in struts has yet to be defined. If the EPIC proteins are involved in interactions with lipids at the epicuticle, they may also play a role in lipid interactions at struts. As *gmap-1* mutants are defective in expansion of the medial layer (Njume et al., 2022), the enlarged EPIC-1 donuts in *gmap-1* might reflect lateral expansion of struts that have not extended vertically.

### Epicuticlins and tandem repeat proteins in the aECM

The epicuticlin family of tandem repeat proteins is conserved throughout nematodes (Betschart et al., 2022). The epicuticlin repeats have two characteristic Y-containing motifs, YGDE and (S/A)GYR. Proteins containing GYR motifs are widespread in arthropod aECMs such as cuticles and eggshells (Cornman, 2010). The significance of the Y-containing motifs remains to be determined but could suggest the epicuticlins are crosslinked by dityrosine crosslinks. Another component of the insoluble cuticle fraction, the tandem repeat protein CUT-2, can form dityrosine crosslinks *in vitro* (Lassandro et al., 1994). Enzymes that catalyze dityrosine crosslinking in nematode cuticle include the Duox dual oxidases (Edens et al., 2001); it will be interesting to examine the role of Duox or other extracellular oxidases in EPIC localization.

Many aECM proteins are composed of imperfect tandem repeats, such elastins (He et al., 2007), gel-forming mucins (Perez-Vilar and Hill, 1999), or silk spidroins (Baker et al., 2022). Epicuticlins are distinctive in containing perfect repeats. For example EPIC-1 repeats 3-6 are perfect, as are EPIC-2 repeats 2 and 7. Other repetitive intrinsically disordered proteins have been identified in *C. elegans* aECMs such as the chitinous pharynx cuticle (Kamal et al., 2022) and suggest that aECMs may involve networks of intrinsically disordered proteins. Relative to imperfect repeats, perfect repeats have a higher tendency to be unstructured (Jorda et al., 2010) and are underrepresented in protein structure databases. The low complexity of such repeats renders them unsuitable for homology-based prediction algorithms such as Alphafold. The tertiary and higher-order structures of epicuticlins are a challenge for future investigation.

Several vertebrate epidermal proteins such as involucrin, filaggrin, trichohyalin, etc are composed of low-complexity tandem repeats and together form the cornified envelope (CE), which replaces the plasma membrane of cornified keratinocytes. Defects in the CE cause a wide range of skin pathologies; for example mutations in filaggrin cause ichthyosis vulgaris and predispose to atopic dermatitis (Sandilands et al., 2009). Involucrin and other scaffold proteins are covalently crosslinked to a monolayer of μ-hydroxyceramides that form the cornified lipid envelope (Jonca and Simon, 2023). Genetic deletion of individual CE components such as involucrin does not have major phenotypic consequences in mice (Djian et al., 2000), although compound mutants can display permeability barrier defects (Sevilla et al., 2007). Such genetic results indicate mammalian epidermal CE components form a robust and redundant network. Analogously, *C. elegans* EPIC proteins do not appear collectively essential for cuticle development or function but may become important in wound repair and potentially other responses to epidermal stress. In conclusion the EPIC proteins define specific compartments in the apical ECM. Despite their lack of highly penetrant developmental phenotypes, they may play roles in stress responses such as wound repair. Our work also underscores the insights that can be gained from analysis of relatively unstudied proteins (Perdigao et al., 2015; Rocha et al., 2023).

## Methods

### General methods

*C. elegans* maintenance followed standard procedures; mutations were confirmed by PCR or sequencing. Strain genotypes, oligonucleotide sequences, and details of new alleles used in this study are listed in Supplemental Table 1.

### Generation of *epic* deletion and knock-in mutations

*epic-1(gk961616)* was generated by the *C. elegans* Million Mutation Project (Thompson et al., 2013) and obtained in strain VC40784 from the CGC. *epic-2(tm7045)* was generated by the Japanese National Bioresource Project and obtained in strain FX7045 from the laboratory of Shohei Mitani. These deletions were outcrossed to N2 2 or 3 times prior to analysis.

We used the melting method (Ghanta and Mello, 2020) to generate larger deletions in *epic-1* and isolated two deletion alleles *ju1930* and *ju1931*; *ju1931* was ∼300 bp smaller than *ju1930* and not analyzed further. To create *epic-2&3* compound mutants we performed CRISPR in the *epic-2(tm7045)* background and isolated three deletions *ju2003, ju2004,* and *ju2005; ju2004* and *ju2005* were 1 bp smaller than *ju2003* and not analyzed in detail.

*epic-1*, *epic-2*, and *epic-3* knock-in strains were generated by SunyBiotech (Fuzhou, China). All knock-ins contain mNeonGreen inserted at the C-terminus with a 3xGAS linker. All three knock-in strains were viable and fertile; as *epic* null mutants are also viable and fertile, we cannot exclude that the knock-in insertions compromise *epic* function in ways we have not detected. EPIC-1::mNG and EPIC-2::mNG KI strains displayed similar expression and localization although EPIC-2::mNG displayed more accumulation in the epidermal secretory pathway. In strain constructions with other cuticle mutants we did not detect enhancement or suppression of Dpy or Bli phenotypes by the *epic* knock-in alleles.

### RT-PCR

Mixed stage animals were collected from large NGM plates, washed 3X with M9 and incubated for half an hour with rotation at RT. Following washing, worm pellets were collected by centrifugation at 2000 *g* and frozen overnight at -80 °C in 1 ml of TRIzol reagent (Invitrogen). RNA was isolated and solubilized in 25 µl DEPC-H_2_O and concentration measured using a spectrophotometer. 10 µg of RNA was treated with 1 µl of DNAse (Invitrogen TURBO kit) at 37 °C for 30 min, purified by Phenol Chloroform method and precipitated using 100% RNA grade ethanol overnight at -20 °C. Final pellets following purification were solubilized in 20 µl of RNAse free water, concentration again measured, and 1 µg of RNA converted to cDNA using ThermoFisher Superscript III RT Kit. The resulting cDNA was used for PCR using cDNA specific primers (see Supplemental Table 1).

### Biochemistry

Biochemical analysis was performed as previously described (Adams et al., 2023).

### Imaging and fluorescence recovery after photobleaching

Widefield fluorescence and DIC imaging were performed on a Zeiss Axioplan M2 imager as described. Conventional confocal imaging and FRAP were performed on a Zeiss LSM800 confocal. Airyscan imaging was performed on a Zeiss LSM900 confocal. 3D SIM was performed on a Deltavision OMX microscope as described, using levamisole immobilization (Adams et al., 2023). TIRF imaging was performed on the OMX SR microscope platform (Cytiva) in TIRF mode using an Olympus ApoN 60x/1.49 objective (APON60XOTIRF). The penetration depth of the evanescent wave was controlled via OMX software (OMX Acquire) and set to maximal stringency, i.e. the shallowest angle prior to loss of signal, yielding z resolutions < 100 nm.

For imaging of EPIC::mNG strains in dauer stage, dauer larvae were generated by starvation. For imaging of adults, animals were aged at least 24 h from mid L4 stage.

HaloTag JF549 ligand staining for visualization of BLI-2::HT in SIM was performed as described (Adams et al., 2023).

EPIC puncta distribution was quantitated from single focal planes of Airyscan processed images of adults 24 h post L4 stage. To measure puncta spacing we drew 10-15 μm line scans along furrow-flanking rows of puncta and counted peaks. To measure puncta density and % area we used a brightness threshold of 56-70 and a particle size range of 0.001-10 μm^2^, in 2-3 ROIs (100-400 μm^2^ each) per image.

### Lipophilic dye staining of epicuticle

Lipophilic dyes were purchased from ThermoFisher. Dye staining followed protocols based on other lipophilic dyes (Schultz and Gumienny, 2012). In brief, healthy unstarved mixed-stage animals were washed into microcentrifuge tubes with M9 and 0.5% Triton X-100, then washed twice with M9. The desired concentration of lipid dye was added to worms in M9 and the animals incubated for 3 h at RT on a rotator. Animals were washed 2-4 times with M9 then allowed to destain on an NGM agar plate for 10-30 min prior to imaging.

For lipid dye staining of mNG-expressing strains we used R18 (Catalog O246) at a concentration of 1 μM, which consistently stained the epicuticle and filled sensory neurons. Under our conditions, R18 stained annuli, furrows, and two to four longitudinal valleys in the adult alae. R18 annular staining was occasionally non-uniform with smaller intense patches of staining (e.g. Figure 6A) or larger less intensely staining patches. At 0.1 μM R18 filled sensory neurons but did not stain the epicuticle; at 10-100 μM R18 stained internal membranes.

For FRAP experiments we used the dialkylaminostyryl dye DiA (4-Di-16-ASP, catalog D3883) at 100 μg/mL. DiA staining was concentrated in furrows and was weaker but visible in annuli; in alae DiA stained 2-4 lines of varying brightness. FRAP was performed using the LSM800 on L4 stage animals.

### Electron microscopy

Larval EM samples were described previously (Witvliet et al., 2021). Epicuticle or plasma membrane thickness was measured using the Plot Profile function in Fiji in line scans perpendicular to the cuticle plane.

### Permeability and wound healing assays

Permeability assays were performed as described (Njume et al., 2022). Needle wounding was performed as described (Xu and Chisholm, 2014) on animals 24 h after the L4 stage.

### Statistical analysis and reproducibility

All statistical analysis used GraphPad Prism 10. All data sets were for normality and parametric or non-parametric tests used accordingly. At least three biological replicates per strain were analyzed independently (e.g. > 3 animals per genotype imaged). Confocal fluorescence images are representative of 5-10 images per condition acquired over at least 3 sessions. In graphs, dot plots show the mean (red line), and SEM (orange error bar).

## Data Availability

Further information and requests for resources should be directed to and will be fulfilled by the lead contact, Andrew Chisholm.

## Acknowledgements

We thank members of the Chisholm and Jin labs for help and discussions and Yishi Jin for comments on the manuscript. We thank Jennifer Adams for initial characterization of *epic* mutant and knock-in strains and Risa Iwazaki (Del Norte High School) for strain construction. We thank Mei Zhen (Lunenfeld Tanenbaum Research Institute) for generous access to developmental EM primary data sets, and Patrick Laurent (Université Libre de Bruxelles) for discussions and reagents. We thank the CGC and the Mitani lab for strains. The *Caenorhabditis* Genetics Center is funded by NIH P40 OD10440. The TEM image of dauer buccal plug is from the WormImage resource of the Center for *C. elegans* Anatomy, funded by NIH R24 OD010943. This work was supported by NIH R35 GM142433 to AME, and NIH R35 GM134970 to ADC.

## Author Contributions

MP designed and performed *C. elegans* molecular genetics, biochemical analyses and confocal microscopy and analyzed data. EMJ performed lipophilic dye staining, FRAP, and HaloTag staining. AME performed 3D SIM and TIRF microscopy. AME and ADC secured funding. MP and ADC wrote the paper with input from other authors.

## Competing Interests Statement

The authors declare no competing interests.

Supplemental Table 1 (Excel file): strain genotypes, oligonucleotide sequences, and allele genotyping information.

## Supplemental Figures

Supplemental Figure 1

EPIC-1::mNG in ringTIRF microscopy of adult cuticle (L4+48 h), 8 successive z sections from external to internal in 125 nm steps. Scales, 5 μm

Supplemental Figure 2

A. Western blot of EPIC::mNG strains and N2 wild type controls. Mixed stage populations were prepared and fractions F1 (cytosolic), F2 (organelle), and F3 (cuticle) isolated (see methods). mNG fluorescence was verified during the cuticle isolation. Western blot probed with anti-mNG antibodies. EPIC::mNG proteins were not detected in F1 or F2 fractions and were detected as multiple high-molecular weight species in the F3 cuticle fractions of knock-in strains. The predicted molecular weights of untagged EPIC proteins after cleavage of the signal sequence are 33.7 kDa for EPIC-1 and 60.6 kDa for EPIC-2; isoelectric points (excluding signal sequences) are 3.99 (EPIC-1) and 4.15 (EPIC-2). B. RT-PCR analysis of *epic* transcripts in mutant strains. The major *epic-1* RT-PCR product is 400 bp, with weaker bands at larger or smaller sizes corresponding to other repeats. The major *epic-2* RT-PCR product is 2 kb, with smaller bands corresponding to internal repeats. C. EPIC-2::mNG localization in adult head cuticle is normal in *epic-1(gk961616)* mutants. Conventional confocal, images 50 x 25 μm. Scale 10 μm.

Supplemental Figure 3

A. Lack of punctate localization of EPIC-1::mNG during L4. Scale, 10 μm. B. Initial stages of EPIC-1 puncta formation in adult. EPIC-1::mNG (cyan) is non-punctate in early young adult (movie frame 2.9). Double label with BLI-1::mSc (magenta). EPIC-1::mNG puncta formation is visible ∼50 min later (movie frame 3.47). Panels are 20 x 20 μm. Scale, 10 μm. C. Quantitation of BLI-1::mSc and EPIC-1::mNG colocalization in ROIs from L4.9-adult, Pearson’s correlation coefficients.

## Notes

### Competing Interest Statement

The authors have declared no competing interest.

### Summary of Updates

Corrected certain panels not showing after PDF conversion of Figures. Corrected citation of epic transcript timing. Corrected Figure 6 to show EPIC-2::mNG not EPIC-1::mNG

