## Supplementary figures and images for "*C. elegans* epicuticlins define specific compartments in the apical extracellular matrix and function in wound repair"

### Supplemental Figures 1-3

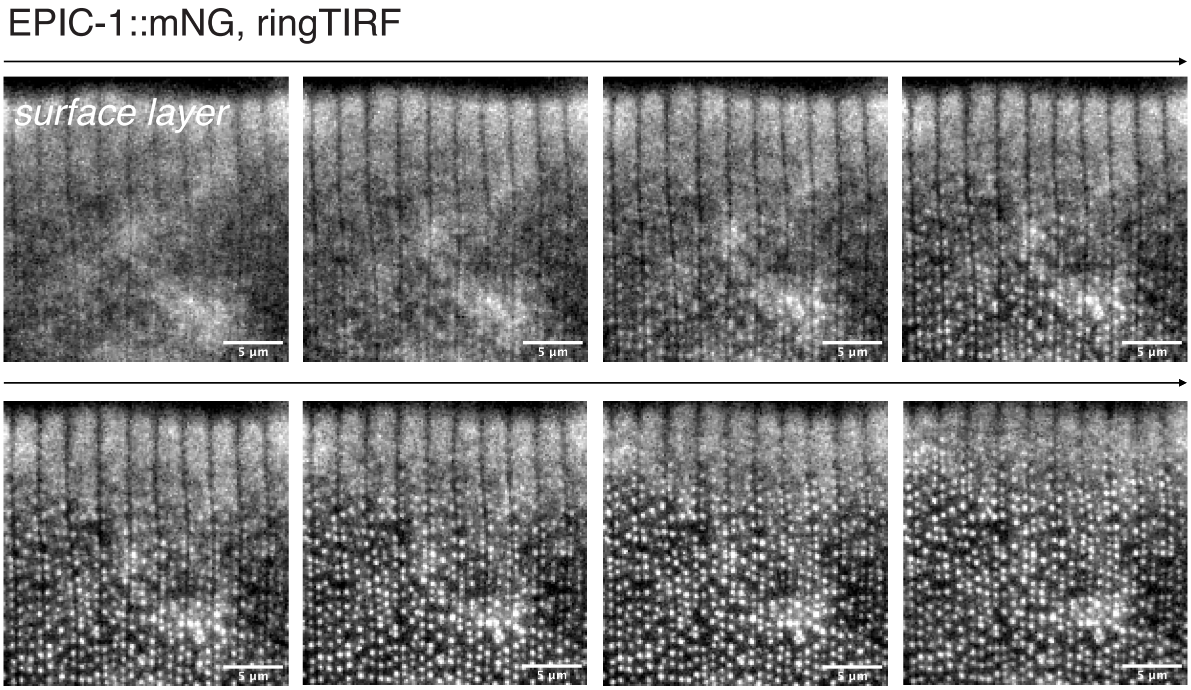


Supplemental Figure 1

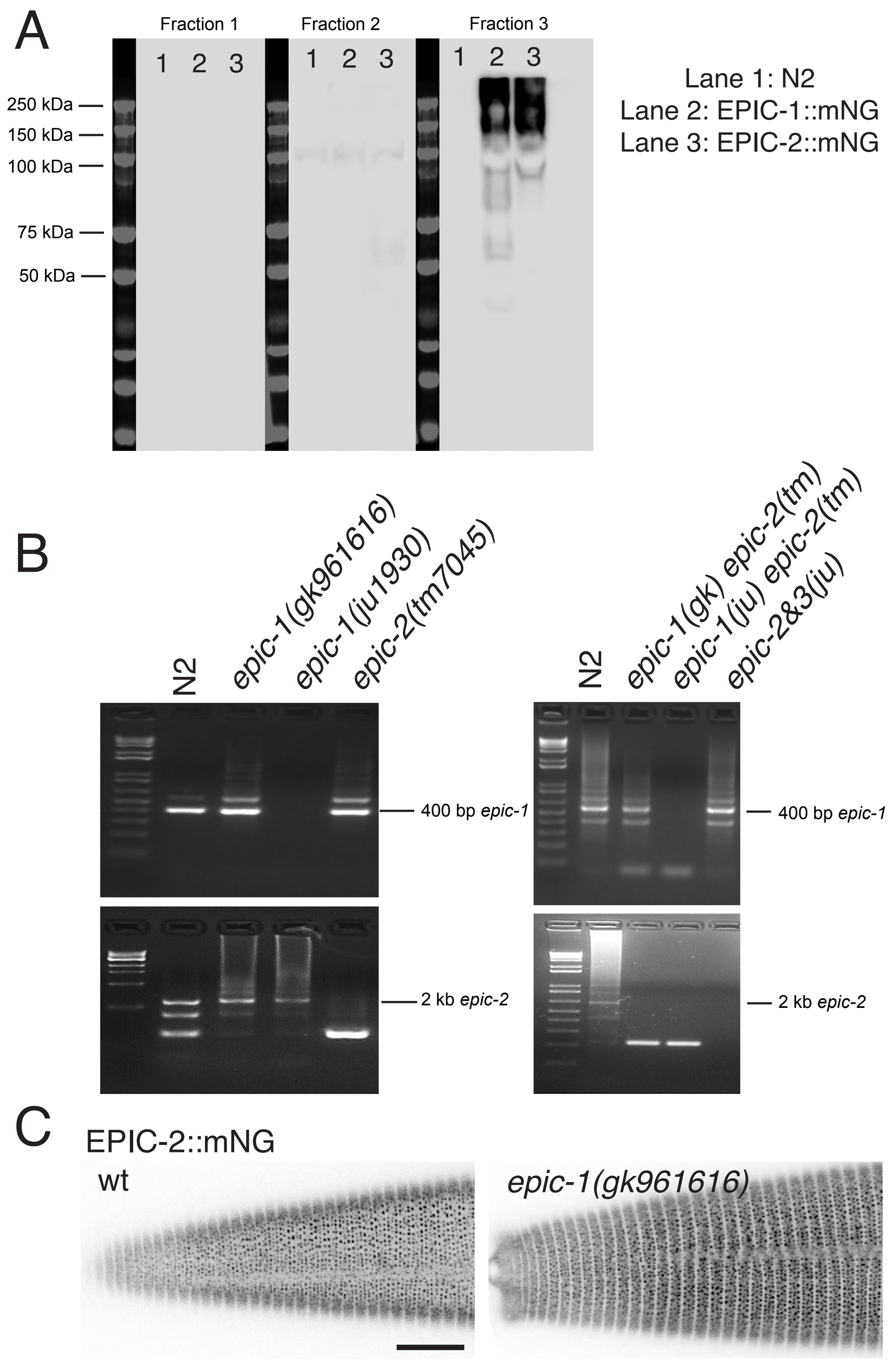


Supplemental Figure 2

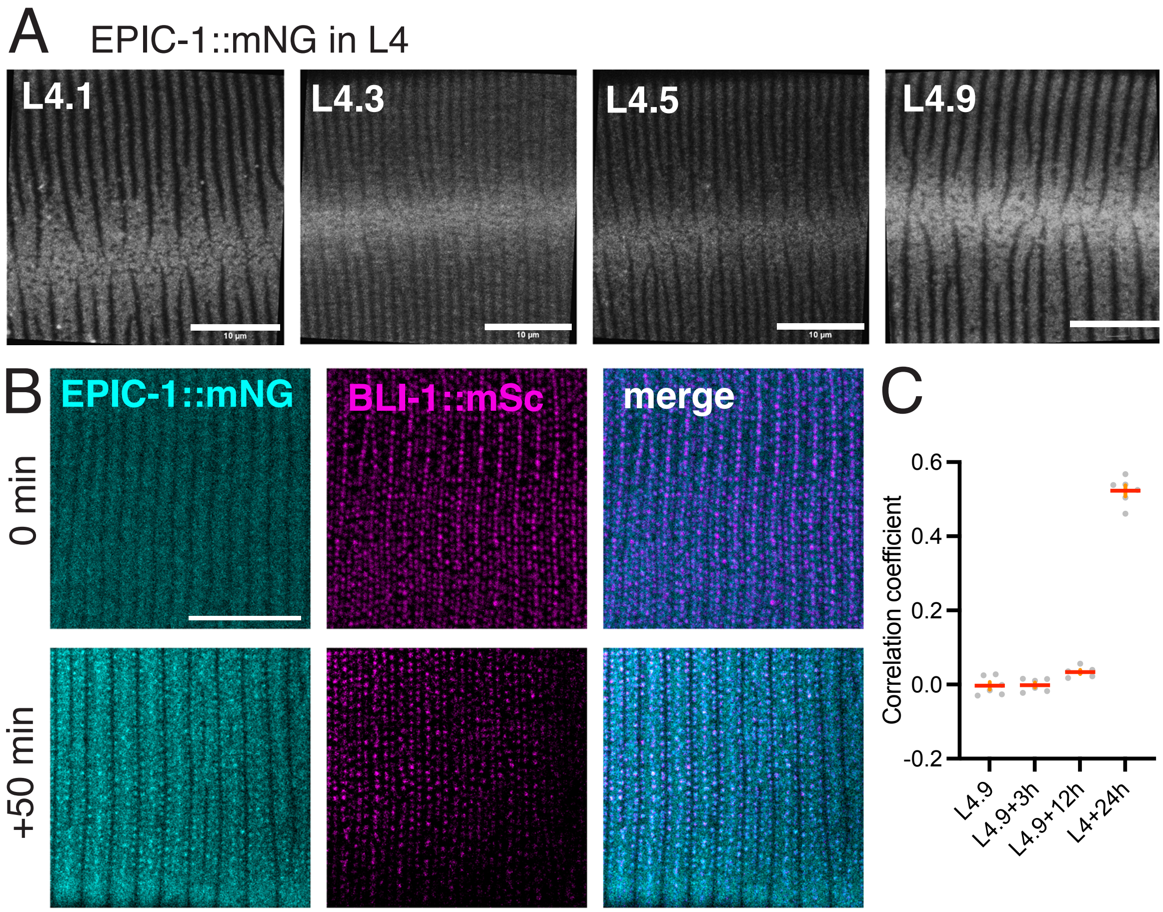


Supplemental Figure 3
